# Ca^2+^-driven PDIA6 phase separation to ensure proinsulin quality control

**DOI:** 10.1101/2024.07.30.605722

**Authors:** Young-Ho Lee, Tomohide Saio, Mai Watabe, Motonori Matsusaki, Shingo Kanemura, Yuxi Lin, Taro Mannen, Tsubura Kuramochi, Katsuya Iuchi, Michiko Tajiri, Kotono Suzuki, Yan Li, Yunseok Heo, Yuka Kamada, Kenta Arai, Mayuko Hashimoto, Satoshi Ninagawa, Yoshikazu Hattori, Hiroyuki Kumeta, Airu Takeuchi, Hiroya Abe, Eiichiro Mori, Takahiro Muraoka, Tsukasa Okiyoneda, Satoko Akashi, Michele Vendruscolo, Kenji Inaba, Masaki Okumura

**Affiliations:** Frontier Research Institute for Interdisciplinary Sciences, Tohoku University, Aramaki, Aoba-ku, Sendai, Miyagi 980-8578, Japan; Biopharmaceutical Research Center, Korea Basic Science Institute, 162, Yeongudanji-ro, Ochang-eup, Cheongwon-gu, Cheongju-si 28119, Korea; Bio-Analytical Science, University of Science and Technology, 217, Gajeong-ro, Yuseong-gu, Daejeon 34113, Korea; Graduate School of Analytical Science and Technology, Chungnam National University, 99, Daehak-ro, Yuseong-gu, Daejeon 34134, Korea; Department of Systems Biotechnology, Chung-Ang University, Anseong, Gyeonggi 17546, Korea; Institute of Advanced Medical Sciences, Tokushima University, Tokushima 770-8503, Japan; Department of Molecular and Chemical Life Sciences, Graduate School of Life Sciences, Tohoku University, Sendai, Miyagi, 980-8577, Japan; College of Life Sciences, Ritsumeikan University, Kusatsu 525-8577, Japan; Department of Materials and Life Science, Faculty of Science and Technology, Seikei University, 3-3-1 Kichijojikitamachi, Musashino-shi, Tokyo, 180-8633, Japan; Graduate School of Medical Life Science, Yokohama City University, Yokohama, Kanagawa 230-0045, Japan; Research Institute of Biomedical and Health Science, Konkuk University, Chungju, Chungbuk 27478, Korea; Department of Biomedical Sciences, School of Biological and Environmental Sciences, Kwansei Gakuin University, Sanda 669-1330, Hyogo, Japan; Department of Chemistry, School of Science, Tokai University, Kitakaname, Hiratsuka-shi, Kanagawa 259-1292, Japan; Biosignal Research Center KOBE University, Rokkodai-cho1-1 Nada-ku, Kobe 657-8501, Japan; Department of Bioresource Science, Graduate School of Agricultural Science, Kobe University, Kobe, 1-1, Rokkodai-cho, Nada-ku, Kobe 657-8501, Japan; Faculty of Advanced Life Science, Global Institution for Collaborative Research and Education, Hokkaido University, Kita 21 Nishi 11, Kita, Sapporo 0110021, Japan; Department of Future Basic Medicine, Nara Medical University, Kashihara, Japan; Department of Applied Chemistry, Graduate School of Engineering, Tokyo University of Agriculture and Technology, Koganei, Tokyo 184-8588, Japan; Department of Chemistry, Centre for Misfolding Disease, University of Cambridge, Cambridge, CB2 1EW, UK; Medical Institute of Bioregulation, Kyushu University, Fukuoka, 812-8582, Japan; Core Research for Evolutional Science and Technology (CREST), Japan Agency for Medical Research and Development (AMED), Tokyo, Japan

## Abstract

The endoplasmic reticulum (ER) plays key roles in protein quality control^1,2^ and dynamic Ca^2+^ storage^3,4^ in eukaryotic cells. However, the protein homeostasis (proteostasis) system that regulates these ER functions is still incompletely characterised. Previous study revealed the importance of oligomerization in the function PDIA1, an ER-resident disulfide isomerase and molecular chaperone, regulates oligomeric states in accordance with client folding^5^. This result suggests that at least some of the 20 members of other PDI family may undergo regulated self-assembly in order to optimally function. Here, we show that Ca^2+^ triggers the phase separation of PDIA6 into liquid-like condensates. In contrast to the condensation mechanism observed for proteins containing low-complexity domains, our results indicate that transient but specific electrostatic interactions occur between the first and the third folded thioredoxin-like domains of PDIA6. We further show that the Ca^2+^-driven condensation of PDIA6 recruits PDIA3 and proinsulin, thus increasing their local concentrations. This process results in the 30-fold enhancement of proinsulin folding and in the inhibition of proinsulin aggregation. Our findings shed light on a condensation-driven Ca^2+^-mediated proteostasis cascade in the ER by revealing how the efficiency of the protein folding process can be enhanced within quality control granules.

## Main

Cells have evolved several strategies to isolate biochemical/biophysical reactions through membrane-bound or membraneless compartmentalization^6,7^. Nearly one-third of nascent polypeptide chains enter the endoplasmic reticulum (ER), in which a proteostasis network, including molecular chaperones and oxidoreductases, guides a wide range of clients to correctly fold. During this process, the proteostasis network prevents aggregation and promotes posttranslational modifications, such as disulfide bond and glycosylation^8^. The ER is also involved in dynamic storage of Ca^2+^, a second messenger in cells, and thus alterations in Ca^2+^ concentrations in the ER cause ER stress, which ultimately leads to apoptosis^9^. The functions of many ER-resident chaperones and oxidoreductases, such as the PDI family, are modulated by continuous fluctuations in the concentration of Ca^2+^ within the ER, which prevents the aberrant accumulation of misfolded or mislocalised polypeptides^10,11^. Ca^2+^-driven proteostasis in the ER, however, remains incompletely understood.

The members of the PDI family are key components of the ER proteostasis system, and they are composed of multiple thioredoxin-like domains and are highly plastic^5,11,12^, resulting in functional diversity. The mechanisms by which they carry out their functions are likely to involve different self-assembly states. For example, PDIA1 monomers form a reversible oligomeric state upon recruitment of unfolded clients, thereby creating a transient catalytic reaction environment in the cavity formed by oligomerization^5^. These structural characteristics suggest that PDI family members may also undergo liquid-like phase separation, which may be relevant for their function. We therefore set out to determine whether PDI family members undergo this process to promote efficient folding in response to Ca^2+^.

## Results

### Ca^2+^ induces the phase separation of PDIA6

To determine whether PDI family members can self-assemble in a Ca^2+^-driven process, we measured their diameters under increasing Ca^2+^ concentrations by dynamic light scattering (DLS) (**Fig. 1a**). Although almost no changes were detectable in the average diameter of highly purified PDIA1, PDIA3, PDIA4, PDIA10, and PDIA15, a Ca^2+^ concentration-dependent increase in average diameter was observed for PDIA6 (**Fig. 1b** and **Extended Data Fig. 1a**). Upon the addition of EDTA, the average diameter of oligomeric PDIA6 returned to its dimeric value (**Extended Data Fig. 1b**) with a reversible catalytic folding activity. At a PDIA6 concentration of >5 μM in the presence of >2.5 mM Ca^2+^, differential interference contrast (DIC) microscopy revealed that numerous spherical droplets were formed. These droplets were approximately several μm up to 50 μm in diameter and induced Ca^2+^-driven droplet formation from the bulk solution (**Fig. 1c, d** and **Extended Data Fig. 1c**). Based on fluorescence recovery experiments after photobleaching (FRAP), fluorescence was recovered in droplets containing mCherry-PDIA6, with a recovery half-time of about 19 sec after photobleaching (**Fig. 1e**). 3D holographic imaging of the refractive index (RI) demonstrated that smaller droplets coalesced together to form larger spherical droplets over time (**Supp. Movies 1 and 2**)^13^, while both the liquid-like droplet fusion and droplet sizes were inhibited by 10 mM NaCl (**Extended Data Fig. 1d**). Droplet formation was also most efficient below pH 7.4 and impaired under relatively high pH conditions (>7.6) (**Extended Data Fig. 1e**), in accordance with the physiological pH condition in the ER. Thus, PDIA6 droplets grow to relatively larger sizes (such as 50 μm diameter) *in vitro* in the presence of Ca^2+^, although the size of droplets in the cells is likely to be regulated by the pH and salt concentration.

**Figure 1.**
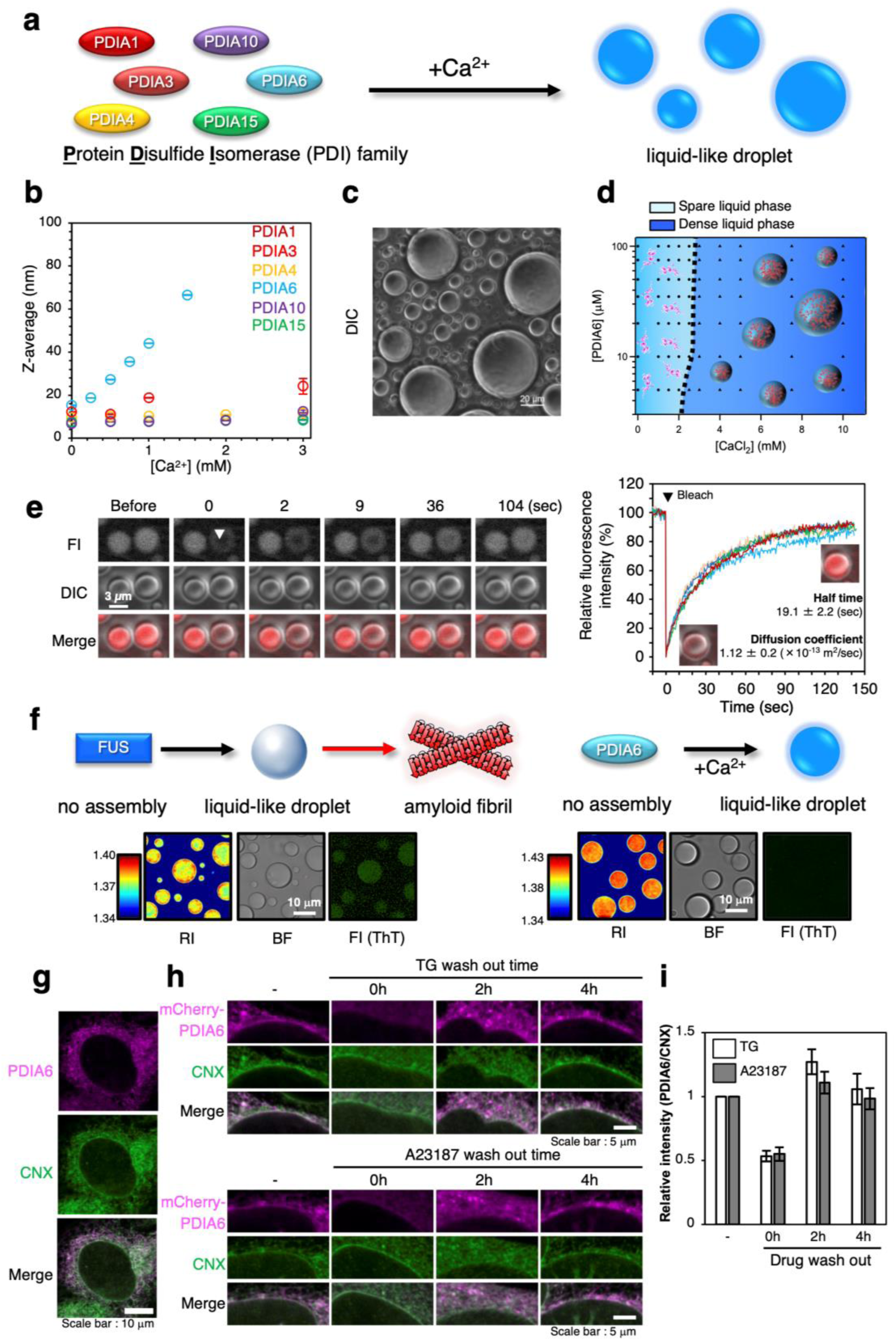
Ca^2+^-driven PDIA6 phase separation. **a**, Schematic representation of the Ca^2+^-induced phase separation of selected PDI family members. **b**, Z-average (the mean diameter of ensemble particles in solution) of PDIA1, PDIA3, PDIA4, PDIA6, PDIA10 and PDIA15 under various Ca^2+^ concentrations. The values are the means ± SDs of three independent experiments. **c**, PDIA6 phase diagram observed by DIC microscopy when 5-100 μM PDIA6 and 0.5-10 mM CaCl_2_ were mixed at pH 7.2. **d**, Quantification of the phase separation assay showing that the amount of PDIA6 remaining in the supernatant ranged from 0 to 10 mM CaCl_2_. The standard deviations of three replicates are represented as error bars. **e**, Confocal fluorescence images of PDIA6 droplets before and after photobleaching. Scale bar, 3 µm. *Left*, rapid recovery of the fluorescence of PDIA6 containing mCherry-PDIA6 after photobleaching. *Right*, The normalized fluorescence intensity of the 5 replicates represented as lines was the half-time (t_1/2_) of fluorescence recovery, and the apparent diffusion coefficients were calculated from the normalized fluorescence intensity of the 5 replicates represented as lines. **f**, Representative 2D RI distribution and fluorescence of FUS or PDIA6 droplets (green, ThT fluorescence; three independent experiments). **g**, U2OS cells were stained for endogenous PDIA6 and CNX. **h**, U2OS cells stably expressing mCherry-PDIA6 were treated with TG and A23187. After the TG or A23187 washout, the cells were stained for mCherry and CNX. **i**, The signal intensities of mCherry-PDIA6 and CNX were statistically analysed using line scan data (n=6 individual experiments).

We next investigated whether amyloid formation can take place within the liquid-like droplets. This question is relevant since it is well-established that fused in sarcoma (FUS), an intrinsically disordered protein with low-complexity sequences associated with amyotrophic lateral sclerosis (ALS), can form amyloid-like assemblies within liquid-like condensates^14,15^. We found that although liquid-like droplets of both PDIA6 and FUS have heterogeneous internal RIs due to their high fluidity, in contrast to FUS, PDIA6 condensates do not exhibit thioflavin T (ThT) fluorescence after 50 min (**Extended Data Fig. 1f**), which is indicative of the maintenance of a liquid-like nature (**Fig. 1f**).

When the endogenous localization of PDIA6 in U2OS cells was investigated, PDIA6 was found to be colocalized with CNX, an ER marker, with some endogenous PDIA6 forming puncta (**Fig. 1g**). To explore the response in behaviour of PDIA6 caused by Ca^2+^ variations inside the ER, we investigated the effects of thapsigargin (TG) and A23187 in U2OS cells overexpressing mCherry-PDIA6. Staining with Mag-Fluo4 AM, which can detect Ca^2+^ in the ER, revealed that TG and A23187 treatments induced a decreased Ca^2+^ in 1 h, but reinflux was observed after 2 h (**Extended Data Fig. 2a**). In response to Ca^2+^ reinflux into the ER, PDIA6 foci appeared at significant levels at 2 h, and PDIA6 co-localized with CNX (**Fig. 1h, i** and **Extended Data Fig. 2b, c**). Taken together, these results indicated that PDIA6 can reversibly regulate its oligomeric state in response to Ca^2+^ *in vitro* and in cells.

### Biophysical and structural characterization of Ca^2+^ binding to PDIA6

To investigate the thermodynamics of Ca^2+^-driven droplet formation, we measured the thermal responses of PDIA6 to Ca^2+^ at various concentrations of NaCl (0, 50, 100, and 150 mM) using isothermal titration calorimetry (ITC) (**Extended Data Fig. 3**). Ca^2+^ bound to PDIA6 at 0 −150 mM NaCl, and Ca^2+^ binding-induced droplet formation was completely suppressed by relatively higher concentrations (>50 mM) of NaCl (**Fig. 2a** and **Extended Data Figs.1c** and **4**). Thus, thermal responses at 0 mM NaCl include both Ca^2+^ binding to PDIA6 and liquid-like condensation, whereas responses at concentrations higher than 50 mM NaCl (50, 100, and 150 mM) reflect only Ca^2+^ binding to PDIA6. By comparing the thermodynamic parameters directly obtained from ITC at 0 mM NaCl (red circles in **Fig. 2b**) and obtained using extrapolation of data at 50-150 mM NaCl (black circles in **Fig. 2b**), we deduced the net thermodynamic parameters of Ca^2+^-induced PDIA6 liquid-like condensation. All ITC thermograms displayed upwards peaks, indicating that the binding of Ca^2+^ to PDIA6 is an endothermic reaction (**Extended Data Fig. 3**). Increasing the NaCl concentration from 0 to 150 mM reduced the magnitude of endothermic heat. Further quantitative analyses showed that with increasing the NaCl concentration, the unfavourable enthalpy changes (Δ*H > 0*) largely decreased from ∼650 to ∼44 cal/mol, and the favourable entropy changes (*T*Δ*S > 0*) decreased from ∼5,500 to ∼4,000 cal/mol (Extended Data Fig. 3). As a result, the reduced net entropy change favourable for complexation decreased the binding affinity between PDIA6 and Ca^2+^ with increasing the NaCl concentration, despite the energy gains resulting from the decrease in the contribution of unfavorable net enthalpy changes.

**Figure 2.**
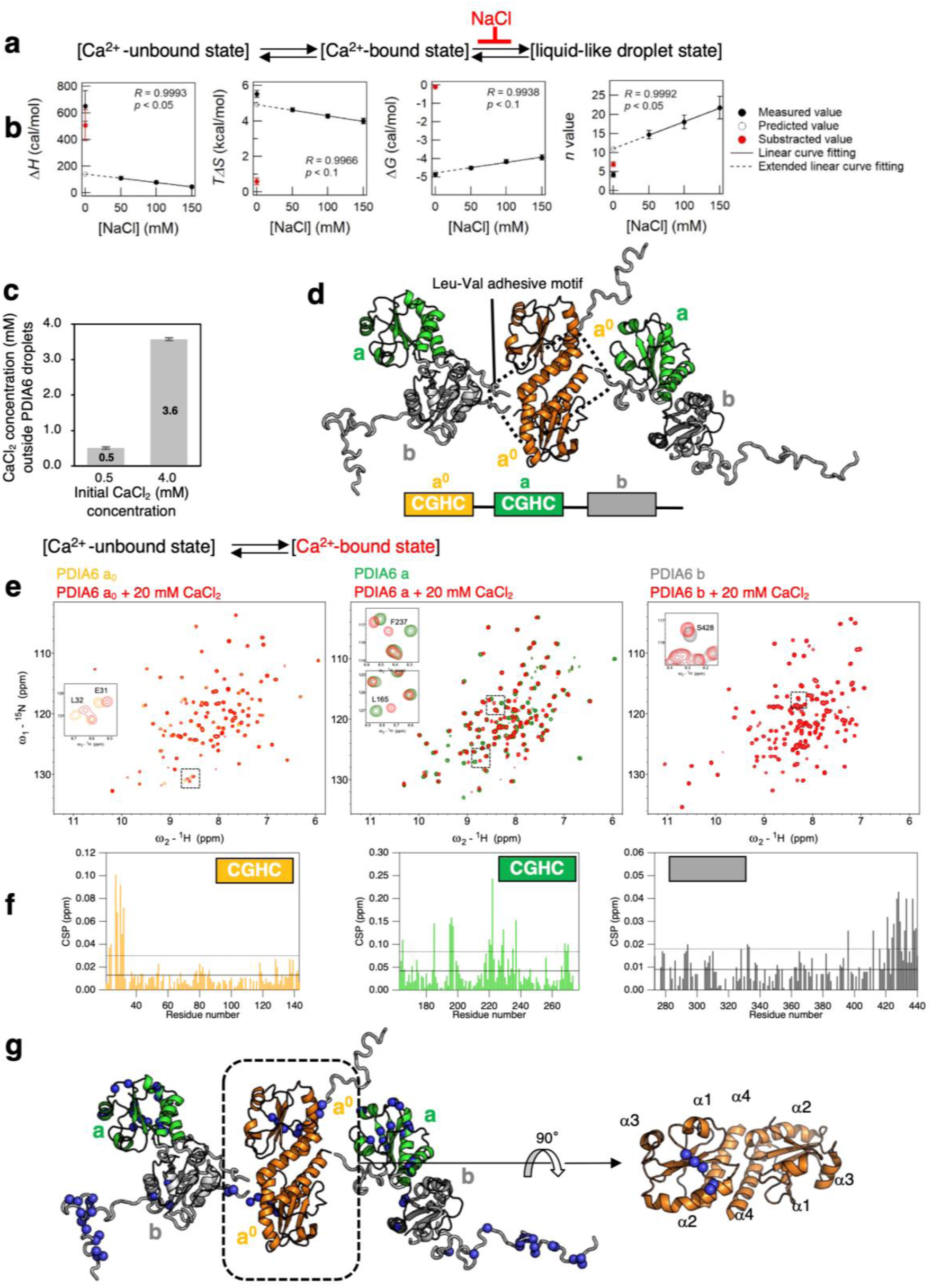
Ca^2+^ binding and Ca^2+^-induced phase separation of PDIA6. **a**, Ca^2+^ binding and Ca^2+^ induced phase separation scheme of PDIA6. **b**, ITC-derived thermodynamic parameters for Ca^2+^ binding to P5 as a function of the NaCl concentration. The solid lines represent the linear fits to the data at 50, 100 and 150 mM NaCl, while the dashed lines are the linear extensions of these solid lines. The black closed circles represent the measured thermodynamic parameters, and the open circles denote the predicted values for 0 mM NaCl based on the dashed line. The red closed circles indicate the differences between the measured data and the predicted data at 0 mM NaCl. The *p* and *R* values for the linear regression analysis are displayed. The error bars show the fitting error. Some error bars are smaller than the symbol size and may not be visible in the figure. **c**, Quantitation of the Ca^2+^ concentration outside PDIA6 droplets. The values are the means ± SDs of three independent experiments. **d**, PDIA6 is composed of two redox-active domains (**a^0^** and **a**) at the N-terminus and a redox-inactive domain **b** at the C-terminus. PDIA6 forms a homodimer via a unique Val-Leu adhesive motif contained in **a^0^**. **e, f and g**, Evaluation of Ca^2+^-binding sites on PDIA6 using NMR. e, ^1^H-^15^N heteronuclear single quantum coherence (HSQC) spectra of the three ^15^N-labelled domains of PDIA6 (left, **a^0^**; middle, **a**; right, **b**) obtained in the absence (orange, **a^0^**; green, **a**; grey, **b**) and presence (red) of 20 mM CaCl_2_. f, Graph representation of the CSPs between the resonances of **a^0^** (left), **a** (middle), and **b** (right). The solid and dotted lines indicate the average and average values added with standard deviation, respectively. g, The CSP mapping to PDIA6 structure. The residues with significant CSPs by Ca^2+^ binding are indicated as blue spheres.

Subsequently, to predict the thermodynamic parameters for Ca^2+^-PDIA6 complexation at 0 mM NaCl without droplet formation, a linear fit was applied to the data at 50, 100, and 150 mM NaCl (**Fig. 2b**). The differences between the measured and predicted values at 0 mM NaCl are attributed to PDIA6 droplet formation. The results showed that PDIA6 droplet formation is an endothermic reaction driven purely by positive entropy changes, Δ*G* = −0.1 ± 0.1 kcal/mol; Δ*H* = 0.5 ± 0.1 kcal/mol; *T*Δ*S* = 0.6 ± 0.2 kcal/mol. This entropic increase can be attributed to the dehydration from PDIA6 molecules induced by liquid-like condensation. This process compensates for the unfavourable entropy loss resulting from the restriction of conformational flexibility and condensation of PDIA6 molecules within the droplets. The extrapolation methodology also revealed the stoichiometry (*n*), which provided information on the number of Ca^2+^ needed to form the droplet. Thus, the stoichiometry of Ca^2+^ was calculated to be ∼7 to form PDIA6 droplets (**Fig. 2b**). When the Ca^2+^ concentration in the supernatant was determined after the PDIA6 droplets were centrifuged, the original Ca^2+^ concentration in the supernatant (4 mM) was decreased to 3.6 mM (**Fig. 1d** and **Fig. 2c**). Consistent with the *n* value obtained by the ITC analysis, the stoichiometry was found to be *n*=8, which was determined by dividing the decreased Ca^2+^ concentration (400 mM) by the protein concentration (50 mM).

PDIA6 is composed of two N-terminal redox-active thioredoxin (Trx)-like domains (**a^0^** and **a**) and a C-terminal redox-inactive Trx-like domain **b**. PDIA6 also forms a homodimer via a unique Val-Leu adhesive motif contained in the **a^0^**, and all domains are located far away, thereby conferring a flexible nature to the protein in solution (**Fig. 2d**)^11^. The sites responsible for Ca^2+^ binding were characterised by nuclear magnetic resonance (NMR) of PDIA6 domains (**Fig. 2e** and **Extended Data Fig. 5**). NMR measurements were performed in the presence of 200 mM NaCl to discriminate between the Ca^2+^-binding and Ca^2+^-induced phase separation. The ^1^H-^15^N correlated NMR spectra of PDIA6 showed Ca^2+^-induced significant perturbations to the resonances from all three domains (**Fig. 2e, f**). Chemical shift perturbation (CSP) mapping revealed that Ca^2+^ binds to multiple specific sites on PDIA6, including the Asp/Glu-rich segment at C-terminus of **b**; the acidic patch in **a^0^** consisting of D27, D28 and E31, and the acidic patch in **a** consisting of E166, E199 and D223. Notably, in the **a^0^** domain, which is essential for dimerization, CSPs were also detected peripheral of α-helix 4, which contains the Leu-Val adhesion motif responsible for dimer formation, suggesting this characteristic motif in the **a^0^** domain is critical for both the Ca^2+^-binding and driving liquid-like condensation (**Fig. 2g**)^11^.

### a^0^ dimerization and a^0^-b interaction are critical for PDIA6 phase separation

To identify the Trx-like domains that are essential for Ca^2+^-induced phase separation, microscopic DIC observation was performed for the individual Trx-like domains. The results showed that slightly smaller droplets were formed only with **a^0^** after Ca^2+^ was added, but almost no droplets were observed with the other individual domains (**Extended Data Fig. 6**). Residual concentration analysis, gel shift assay, and microscopic analyses involving BF and RI were also performed using each of the separated **a^0^**, **a** and **b** domains. One each domain in the presence of Ca^2+^ was added to one another domain and subsequently incubated for 30 min. Then, after centrifugation, the concentration of the supernatant was determined. Notably, only the combination of **a^0^** and **b** decreased the actual measured concentration (red circles in **Fig. 3a**) compared with the theoretical concentration (black circles in **Fig. 3a**), and precipitates were observed after centrifugation. To further detect the decrease in content of each domain, the reaction mixtures were centrifuged, and each domain was quantified by SDS-PAGE (**Extended Data Fig. 7**). An MBP fusion protein was used for **b** since **a** and **b** have almost the same electrophoretic mobility and are difficult to separate. After centrifugation, the remaining concentrations of each domain in the **a^0^**-**a** and **a**-**b** combinations were the same, while the concentrations of each domain in the **a^0^**-**b** combination decreased. Interestingly, when the amount of **b** was excessive and the **a^0^** amount decreased, the **b** amount decreased to approximately 60% (**Extended Data Fig. 7b**). Under the same conditions, the results of each domain titration experiment were examined microscopically, and the droplets were observed only for the **a^0^** and **b** combination, while slightly smaller droplets were detected for the **a** and **b** combination (**Extended Data Fig. 8a**). Furthermore, 3D holographic imaging of the RI revealed that loss of the dimerization motif in the first domain **a^0^** suppressed droplet formation, indicating that the dimerization motif regulates Ca^2+^-induced phase separation (**Fig. 3b** and **Extended Data Fig. 8b**). Consistent with full-length PDIA6 (**Extended Data Fig. 4**), Ca^2+^-induced phase separation between the isolated **a^0^** and **b** domains was suppressed at higher NaCl concentration (**Extended Data Fig. 9**), indicating that transient but specific electrostatic interactions between **a^0^** and **b** domains are essential for Ca^2+^-induced phase separation.

**Figure 3.**
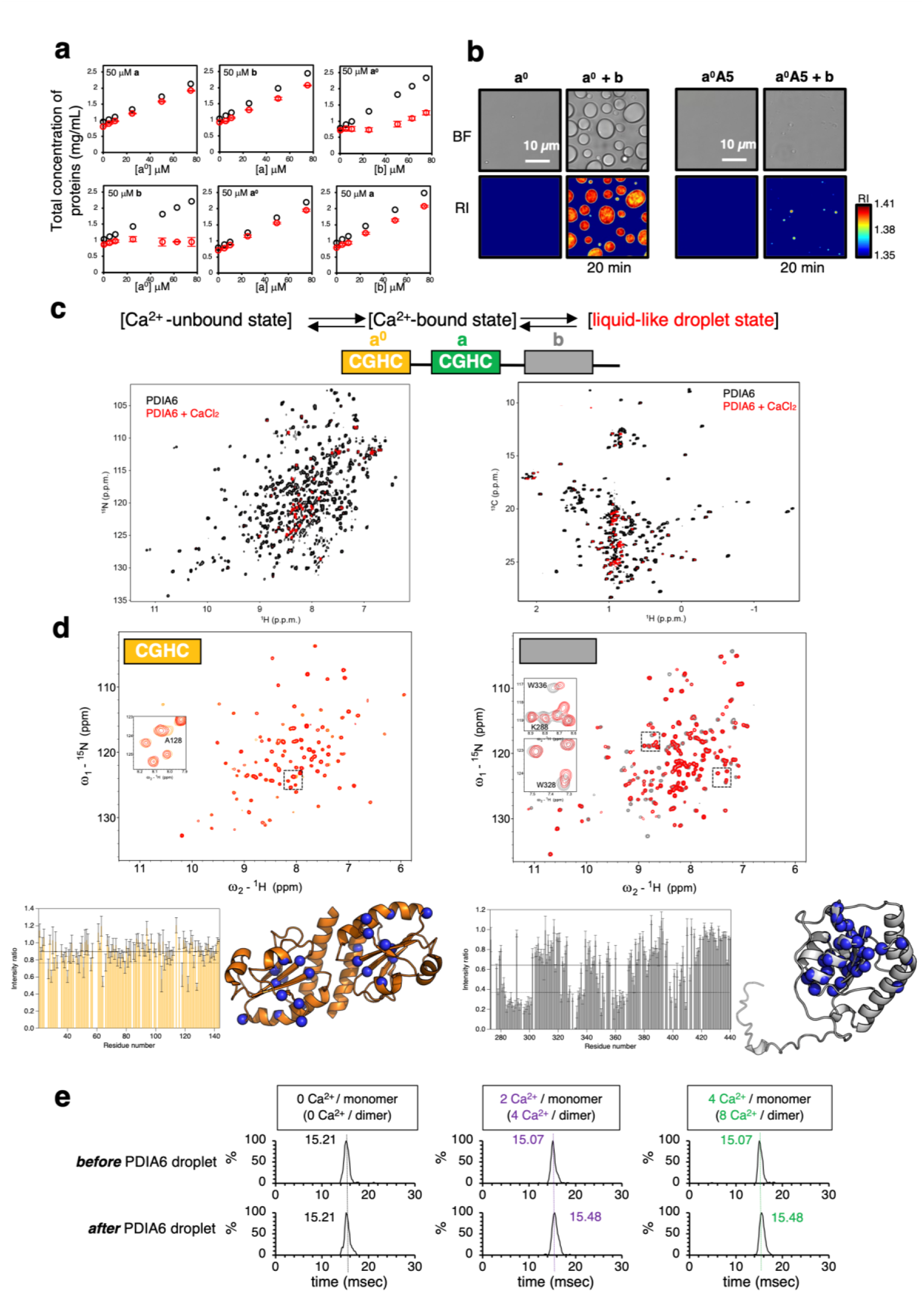
a^0^ dimerization and a^0^-b interactions are critical for the phase separation of PDIA6. **a**, Residual concentrations obtained when each domain was mixed. The black or red open circles indicate the theoretical or measured values, respectively. The values are the means ± SDs of three independent experiments. **b**, Representative 2D RI distribution when **a^0^** and **b** domains (Left) or monomeric mutated **a^0^A5** and **b** (Right) domains are mixed (three independent experiments). **c and d**, NMR investigation of the interaction between PDIA6 molecules during droplet formation. c, The ^1^H-^15^N transverse relaxation optimized spectroscopy (TROSY)-HSQC (left panel) and ^1^H-^13^C heteronuclear multiple quantum coherence (HMQC) (right panel) spectra of [U-^2^H^15^N; Ala-^13^CH_3_; Met-^13^CH_3_; Ile-δ1-^13^CH_3_; Leu, Val-^13^CH_3_/^13^CH_3_]-labelled PDIA6 full length in the absence (black) and presence (red) of 4 mM CaCl_2_. d, ^1^H-^15^N HSQC spectra of ^15^N-labelled **a^0^** (top left) in the absence (orange) and presence (red) of **b** and ^15^N-labelled **b** (top right) in the absence (gray) and presence (red) of **a^0^**. The graphs in the bottom panels show the intensity ratio between the resonances of ^15^N-labelled **a^0^** (bottom left) and **b** (bottom right) in the absence and presence of the other domain. The solid and dotted lines indicate the average and average values, respectively, minus the standard deviation. Residues with large changes in intensity are indicated as blue spheres in the structure of **a^0^** (left) and **b** (right) domains. **e**, Arrival time distributions of PDIA6 19+ charged ions before or after liquid-like condensation observed in the ion mobility MS experiments. The arrival times of 19+ charged ions of 0, 2 and 4 Ca^2+^-bound PDIA6 before or after liquid-like condensation are shown in the plots.

Next, NMR spectra of full length PDIA6 were obtained in the absence and presence of Ca^2+^ to obtain structural information on the PDIA6 droplet. The NMR measurements were performed for the PDIA6 solution without NaCl, in which droplet formation can occur. The addition of Ca^2+^ caused the PDIA6 resonances to significantly broaden, revealing interactions between PDIA6 molecules using multiple areas (**Fig. 3c**). To evaluate the specific interactions between **a^0^** and **b** domains that are essential for Ca^2+^-induced phase separation (**Fig. 3a** and **Extended Data Figs. 8 and 9**), an NMR interaction study was performed for the **a^0^** and **b** domains in the presence of 20 mM CaCl_2_ and 200 mM NaCl. The addition of **a^0^** to isotopically labelled **b** led to significant perturbations in the resonances of **b** domains around the N-terminal α helix (**Fig. 3d**), suggesting that **b** uses a specific region to interact with **a^0^**. On the other hand, the addition of **b** to isotopically labeled **a^0^** resulted in less significant perturbations from sporadic location, suggesting that **a^0^** undergoes multivalent and nonspecific interactions with **b** (**Fig. 3d**).

To detect Ca^2+^-binding or phase separating status by native-MS, ESI mass spectra in 30 or 60 mM NH_4_OAc were measured in the presence of 250 μM CaCl_2_. The droplets were formed in 60 mM NH_4_OAc but impeded in 30 mM NH_4_OAc (**Extended Data Fig. 10a**). Consistent with a previously reported dimerization of PDIA6 in solution^11^, 17+, 18+, 19+ and 20+ ions of PDIA6 homodimer were observed by native-MS (**Extended Data Fig. 10b**). Dimer peaks broadened due to difficulties in desalting and/or desolvation. Furthermore, when coupled with ion mobility mass spectrometry (IM-MS), native-MS can provide insight into conformation of a protein^16^. Dimeric PDIA6 in the 19+ charge state exhibited a shorter arrival time in IM-MS in the Ca^2+^-binding state than in the Ca^2+^-free state (**Fig. 3e**), suggesting the binding of Ca^2+^ to dimeric PDIA6 induced its conformational compaction. On the other hand, PDIA6 under phase separating condition exhibited opposite trend: Ca^2+^-bound dimeric PDIA6 in the 19+ charge state exhibited a longer arrival time in IM-MS than in the Ca^2+^-free state. It is surmised that the PDIA6 homodimer in droplet assumes more extended conformations than those of the dissolved PDIA6 homodimer to achieve weak and multivalent intermolecular interactions between PDIA6s.

### PDIA6 phase separation as an enhanced protein quality control granule

Given that many PDI family members are ubiquitously expressed in the ER, we investigated whether other PDI family members were incorporated into the PDIA6 droplets. A quantitative analysis was performed with the fluorescence intensity from GFP-labelled PDI family members. Consequently, we found that PDIA3 was most efficiently concentrated in PDIA6 droplets (**Fig. 4a** and **Extended Data Fig. 11**). To further clarify the effect of each PDIs on the inside of the droplet, we observed the refractive change inside droplet using 3D holographic imaging of the RI. Consistent with the fluorescence microscopy results (**Fig. 4a**), the RI values of PDIA6 droplets increased to 1.4 in a dose-dependent manner of PDIA3, revealing selective condensation among PDI family (**Fig. 4b**). For PDIA4, which had little effect on condensation according to the fluorescence experiments, the RI values inside the PDIA6 droplets were decreased in manner dependent on the PDIA4 concentration and became sparse even though the size of the droplets did not change significantly (**Fig. 4b** and **Extended Data Fig. 12**).

**Figure 4.**
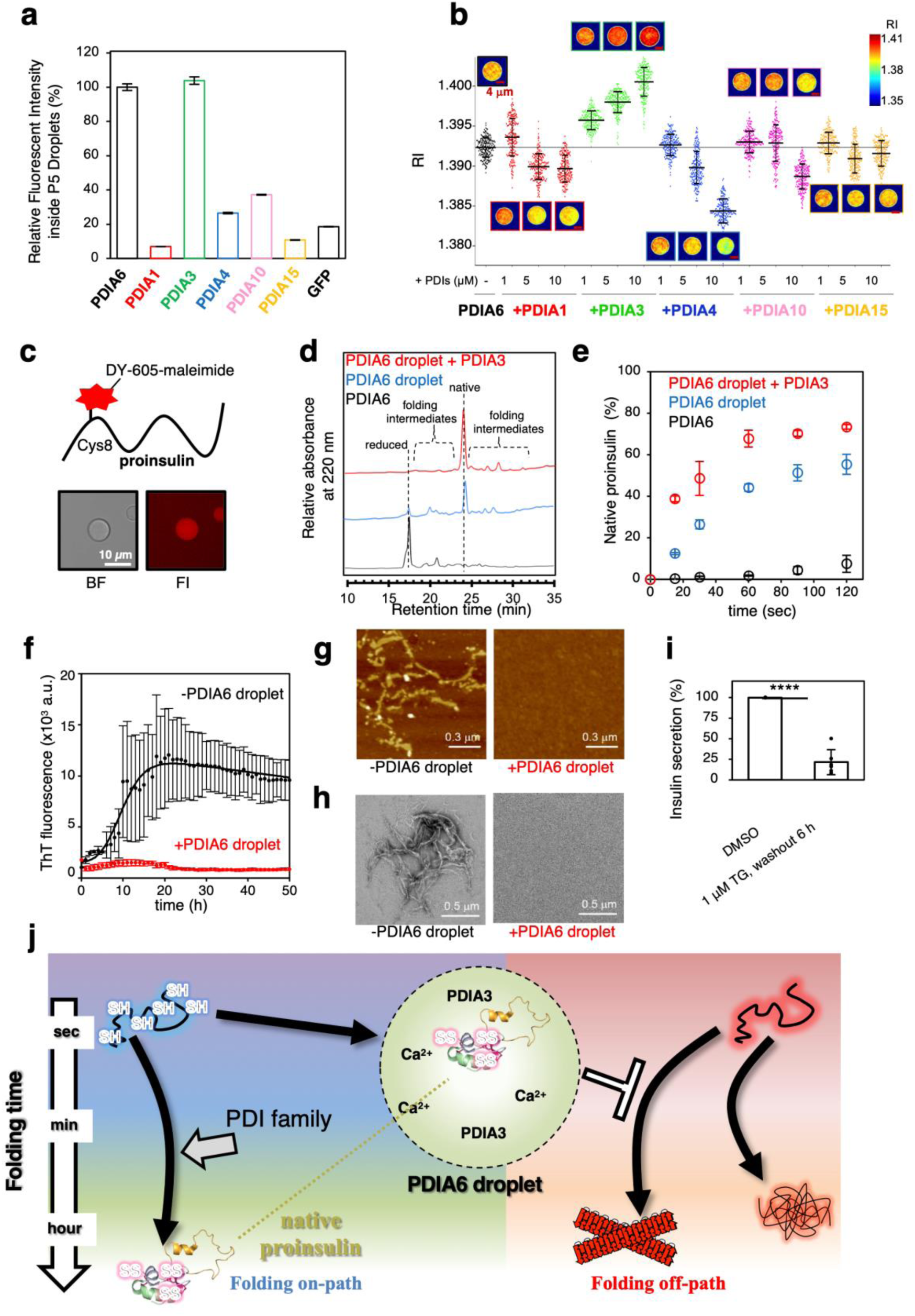
PDIA6 droplets perform proinsulin quality control. **a**, Condensation of the GFP-tagged PDI family into PDIA6 droplets. **b**, RI change plots inside PDIA6 droplets with various PDI family concentrations. Representative 2D RI distributions are indicated. The statistical data were analysed for more than 200 particles in 3 independent experiments. **c**, Condensation of fluorescently labelled proinsulin into PDIA6 droplets. In the labelled proinsulin, 5 Cys residues, except for Cys8, were replaced with Ser, and Cys8 was selectively fluorescently labelled. **d**, Reversed-phase HPLC profiles of the redox state of proinsulin after 60 sec of enzymatic catalysis. **e**, Enzymatic kinetic rates for the proinsulin during oxidative folding (3 independent experiments). **f**, Amyloid fibril formation of proinsulin with all Cys to Ser mutants with and without PDIA6 droplets detected by ThT (3 independent experiments). AFM images (**g**) and TEM images (**h**) with or without PDIA6. **i**, MIN6 cells were treated with 1 µM TG for 16 h. The amount of insulin secreted within 6 h after TG washout was then quantified using ELISA and is expressed as a percentage of the amount secreted by the DMSO-treated control cells. Statistical significance was assessed by a two-tailed paired t test (n = 6). The data are presented as the means ± SDs. ****p < 0.0001. **j**, Proposed model of the PDIA6 phase separation as protein quality control.

Since PDIA6 is known to participate in insulin production and glucose-stimulated insulin secretion within insulin-producing mouse cells^17,18^, the mechanism by which PDIA6 contributes to insulin homeostasis was investigated from the viewpoint of liquid-like PDIA6 droplet. Dye-labelled proinsulin without disulfide bonds was found to be incorporated inside the PDIA6 droplet (**Fig. 4c**), implying that the oxidative folding of proinsulin can take place inside the PDIA6 droplet. We found that 1h was required for PDIA6 in its mono-dispersed state to complete the oxidative folding of proinsulin, and that after 60 sec almost no disulfide bonds were introduced (**Fig. 4d** and **Extended Data Fig. 13**). Notably, the conversion to the native and folding intermediate states of proinsulin occurred within 60 sec after the PDIA6 droplet was added, and the catalysis of oxidative proinsulin folding was further accelerated when the PDIA6 droplet with concentrated PDIA3 was added. The rate constants for disulfide bond introduction (*k*_i_) or native formation (*k*_n_) to proinsulin were determined by single exponential fitting to be *k*_i_PDIA6_ = 0.68×10^-2^ /sec, *k*_i_PDIA6-droplet_ = 5.72×10^-2^ /sec, and *k*_i_PDIA6-droplet-condenced-PDIA3_ = 2.91 /sec, *k*_n_PDIA6_ = 0.15×10^-2^ /sec, *k*_n_PDIA6-droplet_ = 1.81×10^-2^ /sec, and *k*_n_PDIA6-droplet-condenced-PDIA3_ = 4.43×10^-2^ /sec. Thus, the rates of disulfide bond introduction or native formation by PDIA6 droplets and PDIA3-condensed PDIA6 droplets were about 8/426 and 12/28 times, respectively, faster than that by mono-dispersed PDIA6. Since disulfide bonds are not introduced into nascent proinsulin immediately after insertion into the ER, we investigated the effect of PDIA6 droplet on the folding-off pathway of all Cys/Ser mutated proinsulin, which cannot form disulfide bonds. In the absence of PDIA6 droplets, all Cys/Ser-mutated proinsulin are highly prone to aggregation, as monitored by ThT fluorescence, which is indicative of amyloidogenesis. However, PDIA6 droplets suppressed proinsulin aggregation at pH 7.4, a physiological value in the ER (**Fig. 4f**). Atomic force microscopy (AFM) and transition electron microscopy (TEM) images clearly showed proinsulin amyloid fibrils without the PDIA6 droplet (**Fig. 4g, h**).

These results indicate that PDIA6 droplets are capable of chaperone function, i.e. to act as quality control granules, to strongly inhibit protein aggregation and amyloid fibril formation. Ca^2+^-driven PDIA6 droplets incorporate PDIA3 and clients to increase the local concentration of these proteins, resulting in an increase in client folding efficiency. We further estimated the effect of Ca^2+^ depletion in the ER on client folding. When MIN6 cells were treated with TG, the amount of insulin secreted was decreased (**Fig. 4i**), indicating that not only the expression levels of PDIA6^17,18^ but also the concentration of Ca^2+^ in the ER are essential for insulin production. Altogether, Ca^2+^-driven PDIA6 droplets perform protein quality control by enhancing the catalytic activity against the oxidative proinsulin folding, as well as by inhibiting proinsulin amyloid-like aggregation to ensure correct proinsulin folding fidelity.

## Discussion

We reported that PDIA6 undergoes Ca^2+^-induced phase separation and that the resulting condensates act as quality control granules to chaperone the correct folding of proinsulin. Even though the Trx-like domain of PDIA6 is highly soluble, our results suggest that an increase in entropy caused by dehydration in the presence of Ca^2+^ might lead to the liquid-like condensation of Trx-like domains. Unlike the droplet formation mechanism that involves low-complexity domains, transient but specific electrostatic interactions between the first redox-active Trx-like domain and the third redox-inactive Trx-like domain of PDIA6 play a pivotal role in phase separation. Considering the Trx-like domain, almost all PDI family contains at least one Trx-like domain. Most of these domains accommodate a redox-active CxxC motif to chemically catalyse the disulfide introduction/isomerization of clients. On the other hand, at least 13 members of the PDI family contain one or more redox-inactive Trx-like domains, and the **b’** domain of PDIA1 has been extensively studied in relation to client recognition^19^. The current findings indicate that specific interactions occur between redox-inactive **b** and redox-active **a^0^** domain via electrostatic forces, which endows the **b** domain with a physical function for Ca^2+^-driven phase separation. Dimeric **a^0^** then serves as a scaffold for constructing a multivalent intermolecular interaction network with **b** via Ca^2+^. This finding provides important insight into the involvement of redox-inactive Trx-like domains as well as redox-active ones in enzymatic catalysis in the PDI family.

We observe that molecular chaperones have already been reported to regulate liquid-like condensation. For example Hsp70 is incorporated into the nucleolus^20^ and TAR DNA-binding protein droplets^21^ to inhibit solid-like aggregation, and Hsp27 and Kapβ2 also modulate the formation of FUS phase separations^22,23^. Here we reported PDIA6 droplets can selectively recruit folding assistants, such as PDIA3, and greatly accelerate the protein folding reaction. PDIA6 puncta are present in the ER under steady-state conditions, but these puncta are enhanced during stress recovery after Ca^2+^ depletion. PDIA6 is known to inactivate activated IRE1α^24^, an unfolded protein response sensor. Thus, PDIA6 plays a role in restoring the steady state by decreasing the risk of accumulated misfolding clients.

In conclusion, we revealed that PDIA6 is capable of forming droplets through liquid-like condensation, and that such condensates can act as protein quality control granules. The present work contributes to our understanding of the range of Ca^2+^-mediated proteostasis processes based on the liquid-like condensation of PDIs. Thus, our results enrich our knowledge of the proteostasis system by the addition of quality control granules driven by PDIA6. We anticipate that further studies on these quality control granules will lead to a deeper understanding of the molecular origins of ER-related misfolding diseases such as type Ⅱ diabetes.

## METHODS

### Recombinant protein expression and purification

cDNA encoding human PDIs (PDIA1, PDIA3, PDIA4, PDIA6, PDIA10 and PDIA15) without the N-term signal sequence were subcloned and inserted into the NdeI and BamHI sites of the pET15b vector (Novagen, Darmstadt, Germany). The plasmids encoded a 6-His tag at the N-terminus of the proteins. PDIA6 domains (**a^0^**, **a**, **b**) and mutants (**a^0^**-A5, **b**-ND) were constructed using a PrimeSTAR Mutagenesis Basal Kit (Takara Bio, Shiga, Japan). All PDIs used in this study were overexpressed in the *Escherichia coli* strain BL21 (DE3) and purified as described previously^11,25^.

Proinsulin, subcloned and inserted into the pET17b vector (Novagen, Darmstadt, Germany), was overexpressed in the *Escherichia coli* strain BL21(DE3) and purified as described previously^26^.

For the isotopically-labelled proteins used in NMR studies, *Escherichia coli* cells were grown in minimal (M9) medium at 37°C in the presence of ampicillin (100 mg/L). ^13^C/^15^N-labeled protein samples used for resonance assignment were cultured in M9 medium supplemented with ^15^NH_4_Cl (1 g/L; CIL) and ^1^H^13^C-glucose (2 g/L; ISOTEC). The methyl-protonated proteins were prepared as described previously^27,28^. Briefly, for the isotopic labeling of Leu and Val methyl groups, α-ketobutyric acid (50 mg/L) and α-ketoisovaleric acid (85 mg/L) were added to the medium 1 h before IPTG was added. To label the Met and Ala methyl groups, [^13^CH_3_] Met (50 mg/L) and [^2^H,^13^CH_3_] Ala (50 mg/L) were added to the medium 1 h before IPTG was added. M9 medium containing 99.9% ^2^H_2_O were used.

### Dynamic light scattering measurements

PDIA1, PDIA3, PDIA4, PDIA6, PDIA10 or PDIA15 (50 µM) were mixed with 0 to 3 mM CaCl_2_ in 50 mM HEPES-NaOH (pH 7.2) at room temperature. The average Z values of all the mixtures were measured using a Zetasizer Nano (Malvern Instruments, Malvern, UK).

### Quantification of concentration after liquid-like condensation

PDIA6 (5 to 100 µM) was incubated with 0 to 10 mM CaCl_2_ in 50 mM HEPES-NaOH (pH 7.2) at room temperature for 30 min. All the samples were centrifuged at 21,500 × g for 5 min at 25°C, after which the concentration of the supernatant was measured by monitoring the absorbance at 280 nm.

### Refractive index measurements

PDIA6 droplet formation was initiated by adding 4 mM CaCl_2_ to 50 µM PDIA6 in 50 mM HEPES-NaOH (pH 7.2) at room temperature. PDIA6 droplets were observed by a holotomography microscope with a laser-induced fluorescence system (Tomocube, Inc., Daejeon, Korea). The RI and fluorescence images show the XY cross section of the droplet centre. The average RI inside the droplet and their radius were calculated by enclosing the RI image in a circle using TomoStudio version 3.2.8 software (Tomocube, Inc., Daejeon, Korea) and plotted by Igor Pro version 6.36 software (WaveMetrics). The statistical significance of differences was examined by one-way analysis of variance with Tukey honestly significant difference (HSD) post hoc testing. All statistical tests were performed using KaleidaGraph version 4.5.1 software (Synergy Software) at a significance level of α = 0.05

### Residual concentration determination

First, 50 µM PDIA6-**a^0^**, **a** and **b** domains were incubated with 4 mM CaCl_2_ in 50 mM HEPES-NaOH (pH 7.2) at 25 °C. After 10 min, 0, 5, 10, 25, 50, 62.5, 75 or 100 µM PDIA6-**a^0^**, **a**, and **b** domains were added, and the solution was incubated for 30 min at 25 °C. All the samples were centrifuged at 21,500 × g for 5 min at 25 °C. The total concentration of proteins in the supernatant was measured by using a BCA kit (Nakarai Tesque, Kyoto, Japan). For the quantification of each PDIA6 domain in the supernatant, all supernatants were separated by nonreducing SDS‒PAGE. The gel images were captured using a ChemiDoc Touch imaging system (Bio-Rad, Hercules CA, USA), and each band intensity was quantified using ImageJ/Fiji.

### Fluorescence recovery after photobleaching

Droplet formation was triggered by adding 4 mM CaCl_2_ to PDIA6 solutions containing 50 µM PDIA6, 4 µM mCherry-PDIA6, and 50 mM HEPES-NaOH (pH 7.2) at room temperature, and PDIA6 droplets were observed with the 559 nm laser line of a confocal microscope (FV1000, Olympus, Tokyo, Japan). A specific spot in the PDIA6 droplet was bleached at 80% transmission for 0.1 sec, and time-lapse images before and after photobleaching were collected (0.5 s frame rate, 300 frames). The fluorescence intensity of the region of interest was then calculated using FV10-ASW software (Olympus, Tokyo, Japan). The images were processed using ImageJ/Fiji^29^ and GIMP (GNU Image Manipulation Program) software (http://gimp.org). The fluorescence intensity before photobleaching was set to 100%, that immediately after photobleaching was set to 0%, and half the fluorescence recovery and apparent diffusion coefficients were calculated from the normalized fluorescence intensity using Microsoft Excel (Microsoft, Roselle, IL, USA) based on previous studies^30–32^.

### Cell culture

U2OS (ATCC, No. HTB-96) cells were cultured at 37 °C with 5% CO_2_ in Dulbecco’s modified Eagle’s medium (DMEM) supplemented with 10% foetal bovine serum (FBS). The construction of stable cell lines was performed as described previously^33^. First, we generated the doxycycline-induced U2OS/FRT stable cell Line U2OS/FRT/TR, which expresses the tetracycline repressor (TR) from pcDNA6/TR-IRES-puro. The pcDNA6/TR-IRES-puro plasmids were transfected into U2OS/FRT cells using Lipofectamine 3000 (Thermo Fisher Scientific, Waltham MA, USA), the cells^34^ were selected with 2 μg/ml puromycin, and single clones of U2OS/FRT/TR cells were isolated.

### Immunofluorescence

Cells were seeded onto round, 12-mm-diameter coverslips (Matsunami) and fixed with 4% paraformaldehyde in PBS for 10 min at room temperature. The fixed cells were permeabilized with 0.5% Triton X-100 in PBS for 15 min, rinsed, and blocked with 1% bovine serum albumin (BSA) in PBS containing 0.1% Tween-20 (PBST) for 1 h at room temperature. The slides were incubated at 4 °C overnight with primary antibodies (diluted in PBST containing 1% BSA). Primary antibodies against PDIA6 (1:1000; Proteintech, 18233-1-AP), calnexin (1:1000; MBL, M178-3), mCherry (1:1000; Proteintech, 26765-1-AP, 1:1000; Proteintech, 68088-1-Ig) and GRP78 BiP (1:1000; Abcam, ab21685) were used. Unbound antibodies were removed by three 10-min washes with PBST. The slides were then incubated with goat anti-mouse Alexa Fluor Plus 488 (1:1000; Thermo Fisher Scientific, A32723, 1:1000; Thermo Fisher Scientific, A11034) or goat anti-rabbit Alexa Fluor Plus 594 (1:1000; Thermo Fisher Scientific, A32740, 1:1000; Thermo Fisher Scientific, A11032) for 1 h at room temperature, washed, and mounted with Fluoro-KEEPER Antifade Reagent (Nacalai Tesque, Japan). Immunostained cells were examined using a confocal laser scanning microscope (FV1000D; Olympus). Each data series was processed with fixed parameters to enable comparison of the signal intensities.

### NMR measurements

NMR spectra were recorded on a Bruker AVANCE NEO 800 MHz NMR spectrometer equipped with a cryogenic probe (CPTCI(F)), Bruker AVANCE III HD 600 MHz NMR instrument equipped with a TBI probe, Bruker AVANCE III HD 500 MHz NMR instrument equipped with a BBO probe and Agilent UNITY INOVA 600 MHz NMR instrument equipped with a TR5 probe. All NMR spectra were processed using the NMRPipe program^35^ and TopSpin, and data analysis was performed with Olivia (https://github.com/yokochi47/Olivia), SPARKY or POKY^36^.

### Native-MS

ESI-MS and ESI-IM-MS experiments were performed on a SYNAPT G2 HDMS (Waters, Milford, MA) instrument equipped with a nano ESI source with travelling wave ion mobility. The samples were deposited in borosilicate capillaries with an ∼2 µm inner diameter prepared in-house, and a 0.127 mm diameter Pt wire (Sigma‒Aldrich) was placed in the capillary. The experimental parameters for ESI-MS were as follows: capillary voltage, 0.6 kV; source temperature, 70 °C; sampling cone voltage, 10 V; trap collision energy, 20 V; argon gas flow rate, 3.0 mL/min; and backing pressure in the source region, 5.5–5.6 mbar. For ESI-IM-MS, the capillary voltage was set to 0.7 kV, the IM wave velocity was set at 800 m/s, the IM wave height was set at 38 V, and the gas pressure of the IM cell was 2.94-2.92 mbar.

### Uptake assay by fluorescence intensity

Droplet formation was triggered by the addition of 4 mM CaCl_2_ to PDIA6 solutions containing 50 µM PDIA6, 5 µM GFP or GFP-fused PDI family proteins, and 50 mM HEPES-NaOH (pH 7.2) at room temperature, and GFP or GFP-fused PDI family proteins in PDIA6 droplets were observed with the 473 nm laser line of a confocal microscope (FV1200, Olympus). Confocal fluorescence images of the proteins in the PDIA6 droplets were obtained 30–40 min after droplet formation. The images were processed, and fluorescence intensity was extracted using ImageJ/Fiji^29^. The fluorescence intensity of GFP-PDIA6 in PDIA6 droplets was set to 100%, and the relative fluorescence intensity was calculated using Microsoft Excel (Microsoft).

### Insulin ELISA

The insulin secreted from MIN6 cells was quantified using a mouse insulin ELISA Kit (Mercodia 10-1247-01) according to the manufacturer’s protocol. Briefly, MIN6 cells cultured on 24-well plates as described previously (PMID: 33198886) were treated with 1 µM TG (Sigma‒Aldrich, T9033) for 16-24 hours. The cells were then washed three times with serum-free D-MEM and incubated in 0.5 ml of serum-free D-MEM for 6 hours at 37 °C. The cell culture medium was collected and used for ELISA after brief centrifugation to remove detached cells.

### Isothermal titration calorimetry

To determine the binding affinity of Ca^2+^ for PDIA6, ITC experiments were carried out with a VP-ITC instrument (Malvern Panalytical Ltd., Malvern UK). PDIA6 and CaCl_2_ were dissolved in 20 mM HEPES (pH 7.2) containing 0, 50, 100 and 150 mM NaCl. The solutions were degassed for 3 m prior to being loaded into the ITC instrument.

### Thioflavin T fluorescence assay

The kinetics of amyloid formation in proinsulin were monitored using a ThT fluorescence assay at 37 °C. To prepare the stock solution, proinsulin powder was dissolved in 50 mM carbonate bicarbonate buffer (pH 10) with 1 M urea and then centrifuged at 15,000 *× g* at 25 °C to remove insoluble matter. The concentration of proinsulin was determined by UV absorbance at 280 nm with a molar extinction coefficient of 5,560 M^-1^ cm^-1^ and adjusted to a final stock concentration of 60 μM. Subsequently, 3 μM proinsulin solutions in the absence and presence of 50 μM PDIA6 were prepared in 20 mM HEPES (pH 7.2) containing 4 mM CaCl_2_ and 5 μM ThT. The ThT fluorescence assay was conducted using a Synergy H1 microplate reader (BioTek Instruments, Winooski, VT, USA). We loaded each sample (150 μL per well) in triplicate into a 96-well full-area plate (Corning, Inc., Corning, NY, USA) and affixed a sealing film (PowerSeal Cristal View, Greiner-Bio-One, Kremsmünster, Austria) to prevent sample evaporation. All the samples were continuously shaken at 807 cpm with 3 mm stainless steel beads. The fluorescence intensity of ThT was recorded from the top of the plate at excitation and emission wavelengths of 445 and 490 nm, respectively.

The kinetic parameters of amyloid formation were determined by fitting the ThT emission curves using:

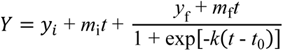

where *y*_i_ + *m*_i_*t* and *y*_f_ + *m*_f_*t* represent the initial and final baseline values, respectively. *t*_0_ denotes the half-life at which the ThT fluorescence intensity reaches 50% of the maximum amplitude, and *k* represents the elongation rate constant. We calculated the lag time using the following relationship: lag time = *t*_0_ − 2(1/*k*). The average and error values of the lag time and elongation rate constant were calculated for three separate samples in a single set.

## Data availability

## Acknowledgment

We are grateful to N. Fukamachi (Tohoku University) and H. Kaneda (Kwansei Gakuin University, Japan) for experimental assistance. We also thank Eunyoung Moon (Korea Basic Science Institute, Korea) for her support with TEM measurements. We are also grateful to Prof. N. Bulleid (University of Glasgow) for deeper discussion.

## Author contributions

Y-HL, TS, and MO designed and supervised this study. MO wrote the first draft. MM, YH, and HK performed the NMR under the supervision of TS. SK and MW performed the HT and biochemical assays under the supervision of MO. IT assisted with the biochemical assays. YLin, YLi, HY, and Y-HL performed the fluorescence assay, ITC, TEM, and AFM under the supervision of Y-HL. TK performed oxidative folding under the supervision of KA and MO. YK performed the insulin secretion assay under the supervision of TO. KIuchi and TM performed the foci formation analysis in cells under the supervision of EM and MO. MM, SK, and SN assisted with the cell biology experiments. MT performed native MS under the supervision of SA. MH and TK prepared the PDIs and several substrates. HA, TM, MV and KI provided expertise in manuscript preparation. All the authors have read and approved the final manuscript.

## Additional information

Reprints and permissions information is available at www.nature.com/reprints.

